# Balanced DNA-to-Cytoplasm Ratio at the 2-Cell Stage Is Critical for Mouse Preimplantation Development

**DOI:** 10.1101/2025.04.30.651589

**Authors:** Tao Pan, Natsumi Taira, Miho Ohsugi

**Affiliations:** Department of Life Sciences, Graduate School of Arts and Sciences, The University of Tokyo, 3-8-1 Komaba, Meguro-ku, Tokyo 153-8902, Japan; Department of Biological Sciences, Graduate School of Science, The University of Tokyo, 7-3-1 Hongo, Bunkyo-ku, Tokyo 113-8654, Japan

**Keywords:** DNA-to-cytoplasm ratio, Preimplantation embryo, Cell size, Ploidy, Mitosis, Cytokinesis

## Abstract

Vertebrate development typically begins with a diploid fertilized egg but can also be initiated parthenogenetically with a haploid maternal genome, though such development fails to reach completion. Among vertebrates, mammalian haploid embryos are particularly prone to arrest during early preimplantation stages for reasons that remain unclear. Here, using mouse embryos with varying cell sizes and genome ploidy, we show that developmental arrest is primarily caused not by haploidy itself but an imbalance between genome ploidy and cytoplasmic volume. Haploid embryos exhibit cytokinesis failure, frequently accompanied by chromosome segregation failure due to malformed spindles beginning at the second mitosis, with about half of the affected blastomeres subsequently arresting before the morula stage. Strikingly, restoring the DNA-to-cytoplasm (D/C) ratio by halving the cytoplasmic volume alleviates these defects. Similarly, diploid embryos with doubled cytoplasmic volume show mitotic and developmental abnormalities resembling those of haploid embryos, whereas tetraploid embryos with the same volume develop normally. These findings demonstrate that a proper D/C ratio, rather than genome ploidy, is critical for mouse embryos to reach the morula stage. Moreover, halving the D/C ratio at later developmental stages, such as the late G2 phase of the 1-cell stage or during the 2-cell stage, progressively improves developmental outcomes. When the D/C ratio is reduced at the end of the 2-cell stage, development proceeds normally. Our findings identify a critical temporal window during which a balanced D/C ratio is essential for mitotic fidelity and successful preimplantation development.

**Graphic Abstract:** 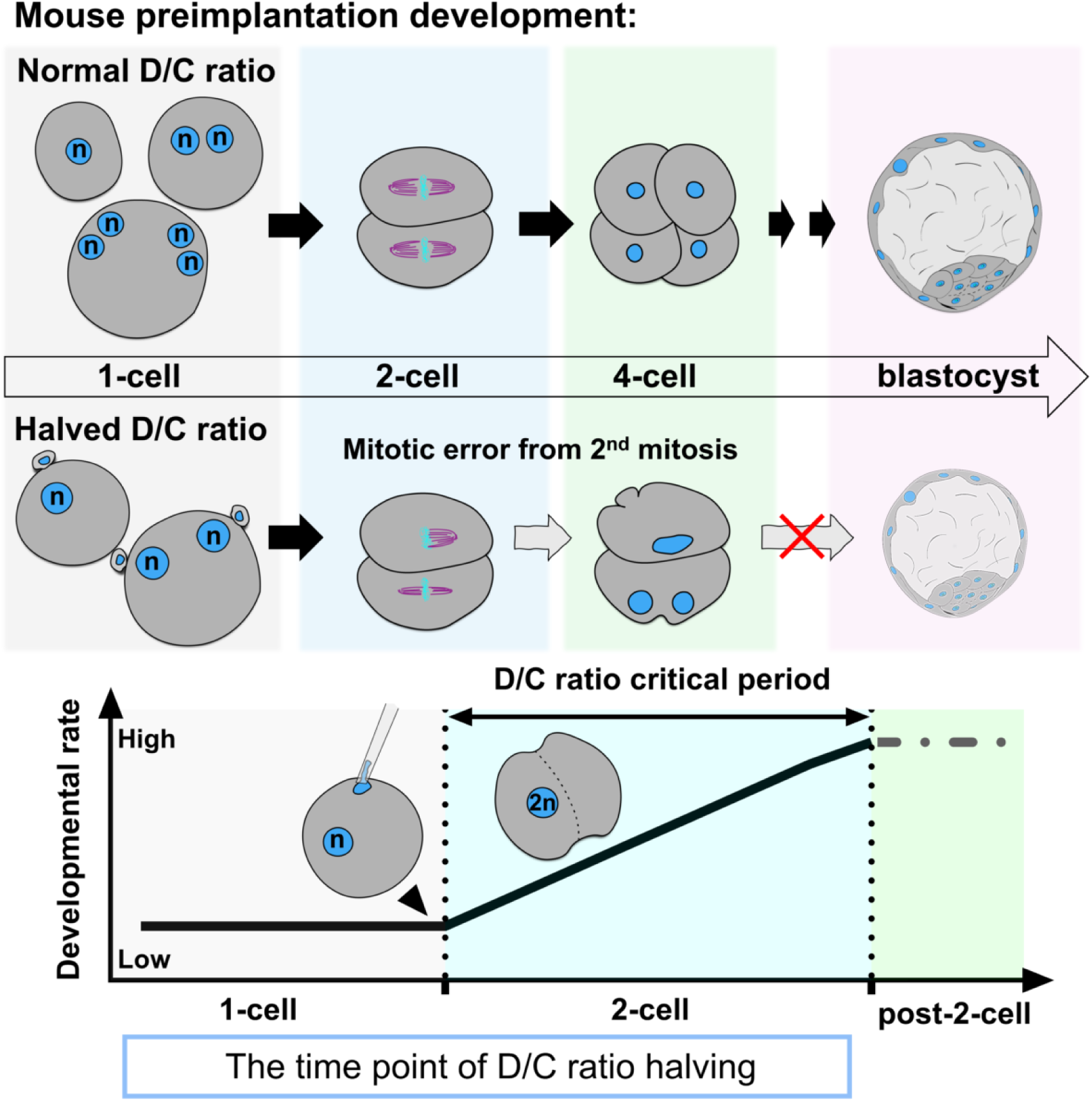

- Halved D/C ratio impairs preimplantation development in mouse embryos
- Halved D/C ratio causes cytokinesis failure from the second mitosis
- Delaying D/C ratio halving to late 2-cell stage alleviates developmental defects

## Introduction

In typical sexual reproduction in animals, embryonic development begins with the fusion of a sperm and an oocyte. Since both gametes complete meiosis before fertilization, the embryo starts with the same ploidy as the parents. However, oocytes can occasionally initiate development without fertilization, either spontaneously or in response to artificial stimulation. These parthenogenetic embryos begin development with half the parental ploidy, though their developmental outcomes vary widely. In Hymenoptera such as bees and ants, fertilized eggs develop into females, while parthenogenetic embryos develop into males, showing that both diploid and haploid embryos can complete development^1^. In contrast, haploid embryos in vertebrates generally arrest during development, although the timing varies among species. Haploid zebrafish show no major defects before organogenesis compared to diploids^2^, and haploid *Xenopus laevis*^3^ and *Xenopus tropicalis*^4^ can progress to the neurula and tailbud stages. In mammals, by contrast, haploid embryos exhibit abnormalities at a much earlier stage. Most arrest before reaching the blastocyst stage, suggesting that developmental failure occurs within five to six cell divisions^5–7^. Since diploid parthenogenetic embryos develop comparably to fertilized embryos^5^, the impaired development of haploid mouse embryos is unlikely to result solely from epigenetic differences between maternal and paternal genomes^8^. Instead, it may reflect mammal-specific regulatory mechanisms that remain unclear.

In addition to genome ploidy, cell size also influences early embryonic development. Since mammalian embryos gain direct access to maternal resources after implantation, oocytes do not require extensive cytoplasmic nutrient storage, resulting in relatively smaller sizes compared to non-mammalian oocytes^9^, typically ranging from 60 to 120 µm in diameter^10^. However, sufficient cytoplasmic volume is still required to support early development. Experimental studies show that artificially reducing mouse oocyte volume significantly impairs post-fertilization developmental potential^11,12^. larger oocytes are often associated with better maturation and higher blastocyst formation rates *in vitro*^13^. However, a larger cell volume is not always advantageous. Increased cytoplasmic volume can dilute checkpoint signals at kinetochores, weakening the spindle assembly checkpoint^12,14^. It can also hinder proper acentrosomal spindle assembly, increasing the risk of chromosome segregation errors and aneuploidy^12^. Although this has not yet been confirmed in preimplantation embryos^15^, early studies in mice showed that artificially enlarging diploid zygotes reduces preimplantation developmental potential^16,17^. These findings suggest that early mammalian embryo size is constrained within an optimal range, but the underlying mechanisms remain unclear.

Among cell size–related factors, the DNA-to-cytoplasm (D/C) ratio has drawn particular attention. It remains relatively stable within specific cell types and is closely linked to biosynthetic capacity and transcriptional regulation, but it can shift under pathological conditions such as cancer and senescence^18–20^. In early embryonic blastomeres, unlike somatic cells, cytoplasmic volume does not increase during cleavage divisions, leading to a progressive rise in the D/C ratio^21^. In fast-developing non-mammalian vertebrates such as *Xenopus* and zebrafish, the D/C ratio contributes to triggering zygotic genome activation (ZGA)^22–25^. Experimental manipulation, either by increasing DNA or reducing cytoplasm, shows that an elevated D/C ratio accelerates ZGA, whereas a reduced ratio delays it^26,27^. These timing changes, however, do not necessarily impair early development. In mice, altering the D/C ratio does not appear to affect the timing of major ZGA ^28^.

In this study, we show that the poor developmental success of parthenogenetic haploid embryos results not only from reduced genome content but also from a halved DNA-to-cytoplasm (D/C) ratio. Restoring the D/C ratio to diploid levels markedly improved their preimplantation development. Conversely, reducing the D/C ratio in diploid embryos significantly impaired development. Stage-specific manipulations further revealed a previously unrecognized regulatory window during the two-cell stage, when a reduced D/C ratio causes mitotic errors and compromises developmental progression.

## Results

### Halving the cytoplasmic volume in haploid embryos improves preimplantation development

Since activated oocytes and early cleavage-stage blastomeres maintain similar sizes regardless of ploidy, the D/C ratio in haploid embryos is about half that of diploids. Thus, when the D/C ratio of a diploid embryo (Dip) is set to 1.0, that of a haploid embryo (Hap) is 0.5, reflecting ploidy-dependent differences in cytoplasmic volume (Fig. S1A-B). We hypothesized that the reduced D/C ratio, rather than haploidy itself, primarily accounts for the lower developmental rate in Hap. To test this, we established a non-invasive method to induce symmetric division at the end of meiosis II, replacing the normal asymmetric division that extrudes the second polar body, to generate haploid embryos with halved cytoplasmic volume. Treating Metaphase-II (Meta-II) oocytes with CK-666, an Arp2/3 complex inhibitor, has been shown to cause spindle relocation from the periphery to the center of the oocyte^29^ (Fig. 1A; Fig. S1C). Sequential treatment with CK- 666 and parthenogenetic activation stimuli yielded nearly symmetric division in ∼30% of activated oocytes, producing half-sized haploid embryos (Halved-Hap) (Fig. 1B).

**Figure 1.**
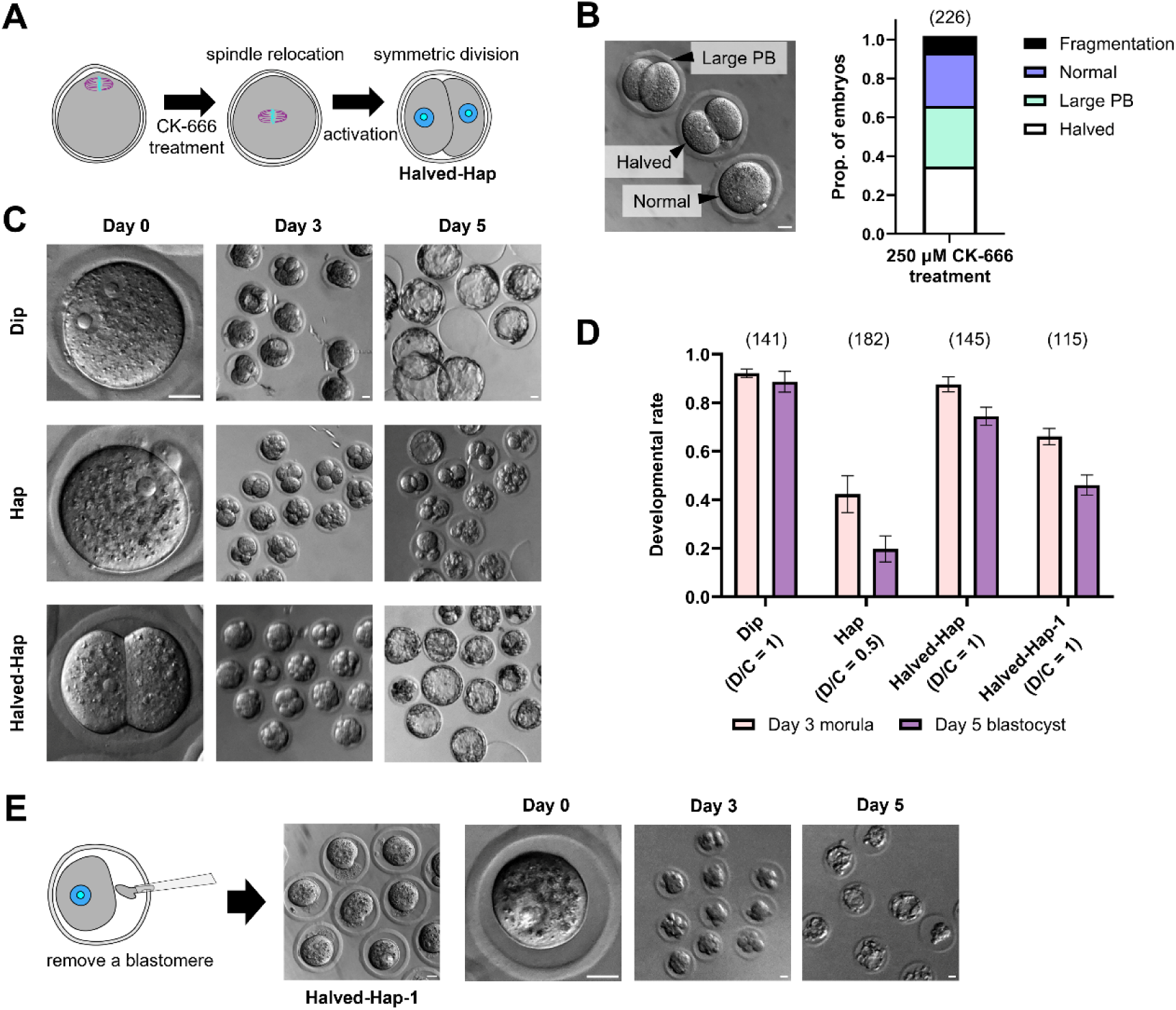
Half-sized haploid embryos exhibit improved preimplantation development compared to normal-sized haploid embryos. (A) Schematic of half-sized haploid embryo (Halved-Hap) production. See also Fig. S1C. (B) Bright-field image and bar graph showing proportions of embryos treated with 250 μM CK-666 (5 experiments). (C) Bright-field images of normal-sized diploid (Dip), haploid (Hap), and Halved-Hap at Days 1, 3, and 5 post-activation. (D) Proportions of embryos reaching the morula (Day 3) and blastocyst (Day 5) stages (mean ± SEM; ≥6 experiments). (E) Representative images of Halved-Hap-1 production and embryos cultured at Days 0, 3, and 5. Scale bars, 20 μm. Sample sizes are shown in parentheses above each bar.

Dip, Hap, and Halved-Hap were cultured for 5 days with imaging every 24 hours, and developmental rates to the morula (∼72 hpa, Day 3) and blastocyst (∼120 hpa, Day 5) stages were quantified (Fig. 1C). Notably, despite being haploid, Halved-Hap showed higher developmental rates, with over 70% reaching the blastocyst stage by Day 5 (Fig. 1D). They began as two equivalent cells, resulting in twice the cell number of Hap, which may confer a developmental advantage. To test this, one blastomere was removed by aspiration to generate embryos hereafter referred to as Halved-Hap-1 (Fig. 1E), which still developed more efficiently than Hap (Fig. 1D–E). Furthermore, fusing the two blastomeres of 2-cell-stage Hap via electrostimulation produced a single blastomere with a diploid genome but a halved D/C ratio relative to 2-cell stage Dip (Fig. S1D). Despite being diploid, these embryos exhibited low developmental efficiency (Fig. S1E–F). Together, these findings suggest that the D/C ratio, rather than ploidy, is the key determinant of preimplantation development.

### Haploid embryos tend to exhibit a prolonged 2-cell stage and mitotic abnormalities at the second and third division

Since Hap exhibited a significantly lower developmental rate before the morula stage compared to Dip, Halved-Hap, and Halved-Hap-1 (Fig. 1D), we next examined abnormalities in Hap that might contribute to developmental arrest. Bright-field microscopy revealed that over 95% of Dip, Hap, and Halved-Hap completed the first cleavage and reached the 2-cell stage by 24 hpa. However, by 48 hpa, more than 30% of Hap remained at the 2- or 3-cell stage, whereas over 90% of Dip and Halved-Hap reached the 4-cell stage (Fig. 2A). Immunofluorescence at 44–48 hpa showed that nearly 40% of the arrested Hap blastomeres contained two or more nuclei (Fig. 2B, C), suggesting increased susceptibility to errors during the second mitosis.

**Figure 2.**
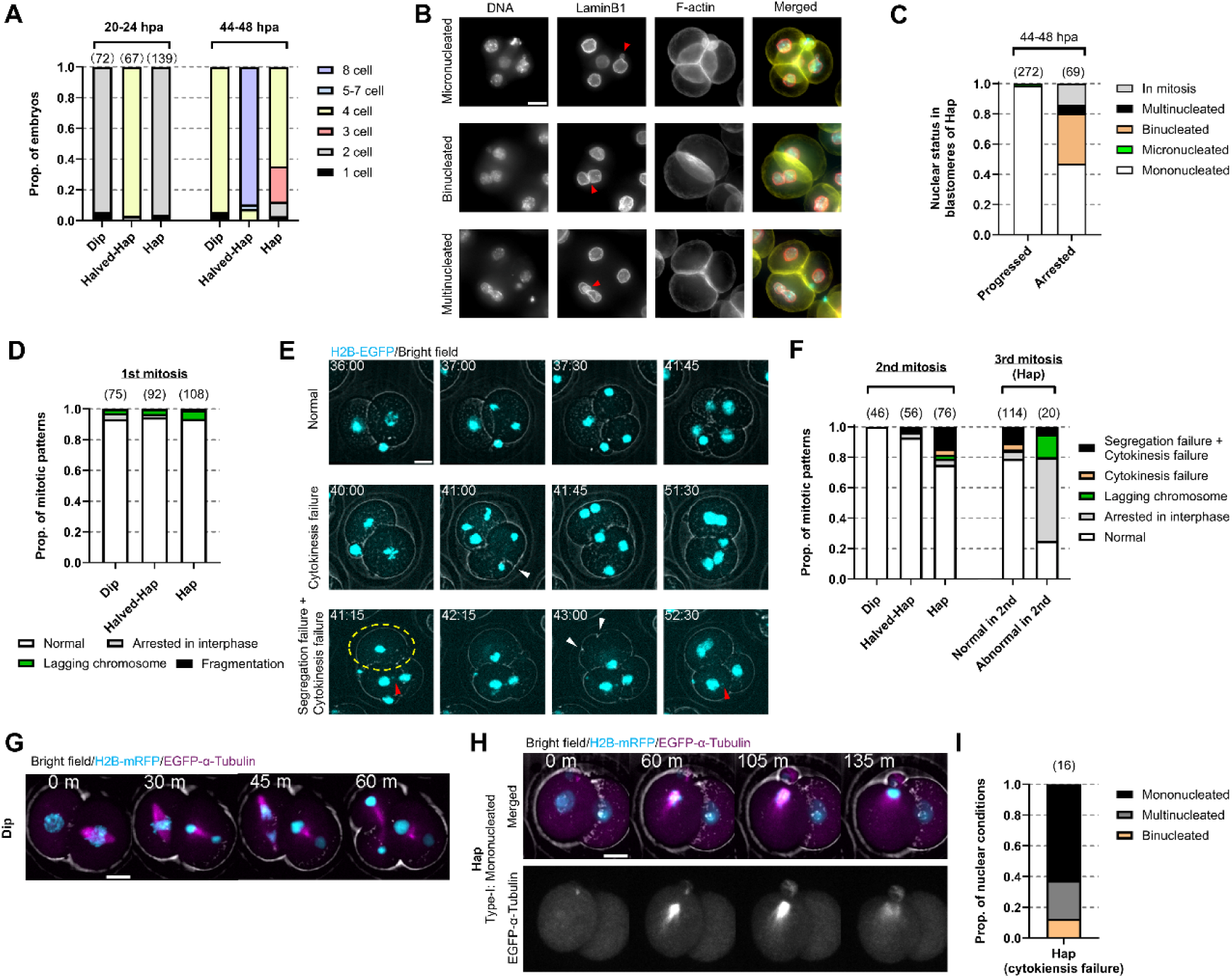
Normal-sized haploid embryos show abnormalities from the second mitosis. (A) Proportions of embryo statuses at 20–24 and 44–48 hours post-activation (hpa). Statuses are represented by cell number rather than developmental stage (≥3 experiments). (B) Z-projected immunofluorescence images of Hap at 44–48 hpa. Red arrows indicate nuclei with abnormal morphologies. In merged images, DNA, Lamin B1, and F-actin are shown in cyan, red, and yellow, respectively. (C) Nuclear status of Hap blastomeres that completed cytokinesis (progressed) or remained 2-cell-sized (arrested) at 44–48 hpa (≥3 experiments). (D) First mitosis outcomes in Dip, Halved-Hap, and Hap (3 experiments). (E) Still images from time-lapse imaging of the second mitosis in Hap expressing histone- H2B-EGFP (cyan, z-projected). Timestamps indicate hpa. White arrows, incomplete cleavage furrows; yellow dashed lines, blastomeres with segregation and cytokinesis failure; red arrows, micronuclei. See also Video S1. (F) Second mitosis outcomes (left) and third mitosis outcomes in Hap based on second mitosis status (right) (≥3 experiments). (G) Still images from time-lapse imaging of the second mitosis in microinjected Dip expressing mRFP1–H2B (cyan, z-projected) and EGFP–α-tubulin (magenta, z-projected). Timestamps indicate time (minutes) since NEBD. See also Video S2. (H) Still images from microinjected Hap showing cytokinesis failure (Type-I, mono- nuclei). Timestamps show time since NEBD. See also Video S2. (I) Nuclear statuses during second mitosis in microinjected Hap blastomeres with cytokinesis failure (3 experiments). Scale bars, 20 μm. Sample sizes in parentheses above each bar.

To further examine cell cycle and mitotic progression, Dip, Hap, and Halved-Hap from histone-H2B-EGFP knock-in oocytes were subjected to time-lapse imaging. Consistent with bright-field observations, over 95% completed the first mitosis (Fig. 2D). Most Hap entered the second mitosis but with several hours of delay compared to Dip and Halved- Hap (Fig. 2F, S2A). Hap also exhibited prolonged mitosis (Fig. S2B), and over 20% of blastomeres showed defects such as severe chromosome segregation errors, including nondisjunction, as well as lagging chromosomes and cytokinesis failure. These abnormalities resulted in either polyploid mononucleated cells or cells containing two or more nuclei. (Fig. 2E-F; Video S1). In cases of cytokinesis failure, a cleavage furrow formed but regressed without completing division (Fig. 2E, white arrows). In contrast, Dip and Halved-Hap showed significantly fewer abnormalities (Fig. 2F). Among Hap blastomeres with second mitosis defects, 55.0% arrested before the third mitosis, 20.0% showed further abnormalities during the third mitosis, and only 25.0% progressed normally (Fig. 2F). Even among cells completing the second mitosis, 21.1% showed defects in the third. These results indicate that reduced D/C ratio compromises mitotic fidelity from the second mitosis onward, leading to cumulative defects and developmental arrest in Hap.

To investigate the cause of cell cycle delay and mitotic abnormalities in Hap, we first examined whether DNA damage was involved. However, γH2AX foci were rarely detected in Hap nuclei during G2 of the 2-cell stage (Fig. S2C, D). We then assessed spindle formation by visualizing microtubules and chromosomes during the second mitosis. In Dip and Halved-Hap, bipolar spindles formed correctly, and chromosome alignment, segregation, and cytokinesis proceeded normally (Fig. 2G, S2E). In contrast, Hap frequently failed cytokinesis and showed three major types of spindle defects with associated chromosome misbehavior. The most common involved monopolar spindles, where chromosomes moved toward a single pole, forming one nucleus (Type-I, Fig. 2H- I; Video S2). Asymmetric bipolar spindles were also observed, often resulting in lagging chromosomes and multinucleated cells (Type-II, Fig. S2F). In some cases, bipolar spindles formed, and segregation appeared normal, but cytokinesis still failed; these spindles tended to be short and occasionally bent (Type-III, Fig. S2G). These findings imply that defective spindle assembly is one of the causes of abnormal chromosome dynamics and cytokinesis failure, thereby contributing to developmental arrest in Hap.

### Halving the D/C ratio of diploid embryos by doubling the cytoplasmic volume induces developmental defects similar to haploid embryos

To further assess the importance of the D/C ratio, we generated diploid and tetraploid embryos with doubled cytoplasmic volume by electrofusing two Meta-II oocytes (Fig. 3A, S3A). Electrostimulation induced simultaneous fusion and activation, resulting in diploid embryos with two second polar bodies and a relative D/C ratio of 0.5 (Doubled- Dip). When cytochalasin B was added to inhibit polar body extrusion, tetraploid embryos were generated with the same cytoplasmic volume and a relative D/C ratio of 1.0 (Doubled-Tet) (Fig. 3A, S1A-B).

**Figure 3.**
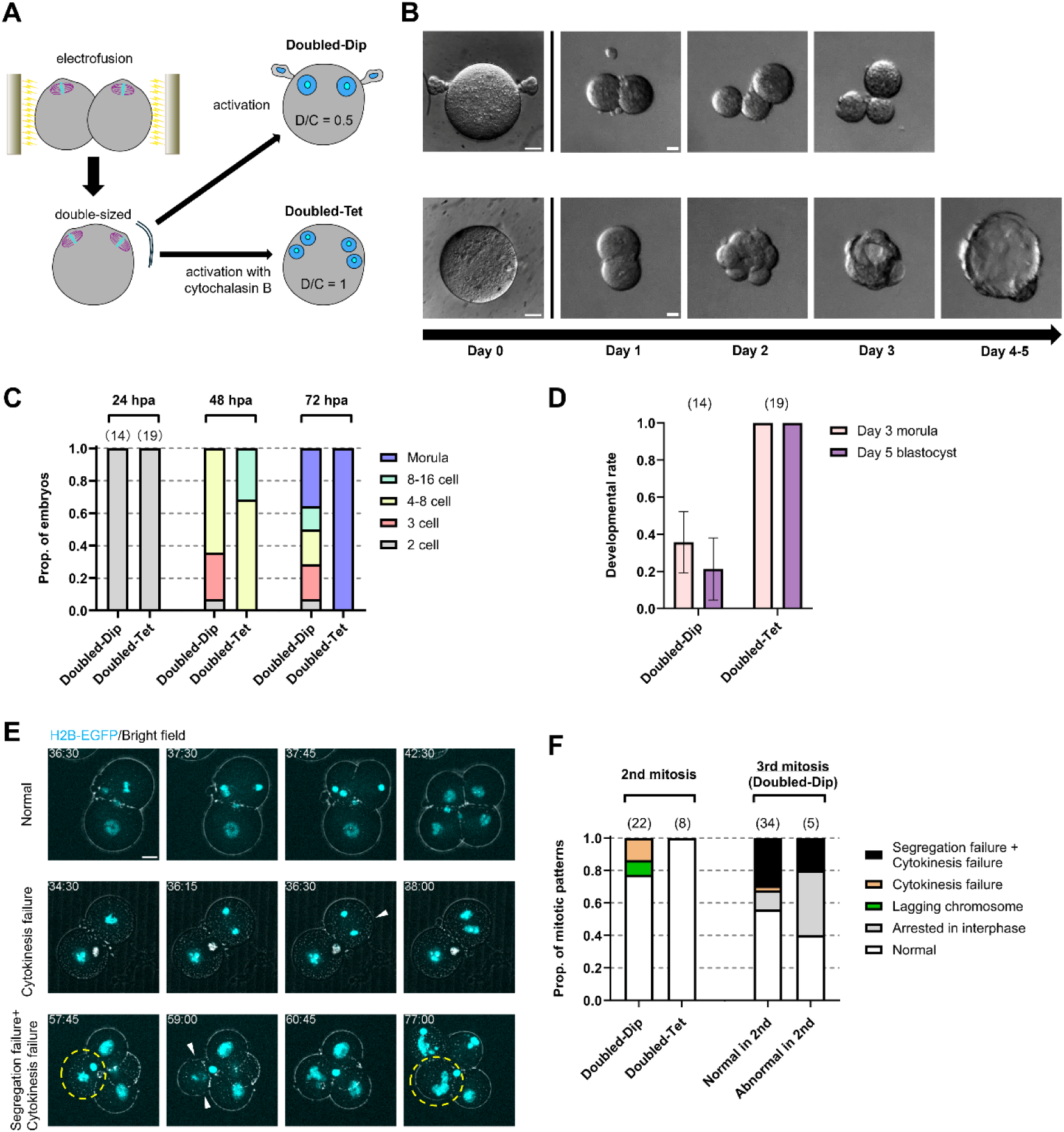
Doubled-sized diploid embryos show developmental defects similar to normal-sized haploid. (A) Schematic of double-sized diploid (Doubled-Dip) and tetraploid (Doubled-Tet) embryo production. (B) Bright-field images of Doubled-Dip (top) and Doubled-Tet (bottom) undergoing symmetric division on Day 1. See Fig. S3C-D for asymmetrically divided Doubled-Tet. (C) Proportions of embryo statuses at 24, 48, and 72 hpa (≥3 experiments). (D) Proportions of embryos reaching morula (Day 3) and blastocyst (Day 5) stages (mean ± SEM; ≥4 experiments). (E) Still images from time-lapse imaging of the second mitosis in Doubled-Dip expressing histone-H2B-EGFP (cyan, z-projected). Timestamps indicate hpa. White arrows, incomplete cleavage furrows; yellow dashed lines, Segregation failure + Cytokinesis failure blastomeres; red arrows, micronuclei. See also Video S3. (F) Second mitosis outcomes in blastomeres from both embryo types (left), and third mitosis outcomes in Doubled-Dip based on second mitosis status (right) (≥5 experiments). Scale bars, 20 μm. Sample sizes in parentheses above each bar.

As expected, about 40% of Doubled-Dip, despite completing the first division, arrested at the 2- or 3-cell stage, and only 35.7% and 21.4% reached the morula and blastocyst stages, respectively (Fig. 3B–C). Immunofluorescence revealed frequent binucleated blastomeres in embryos arrested before the 8-cell stage, similar to Hap (Fig. S3B). In contrast, nearly half of the Doubled-Tet showed asymmetric or abnormal cleavage during the first division, likely due to failure in the assembly of a single bipolar spindle in the presence of four pronuclei and a large cytoplasmic volume (Fig. S3C). Despite irregular blastomere composition, most developed to the blastocyst stage (Fig. S3D). Since the D/C ratio after Day 1 was not quantifiable, we analyzed embryos that underwent symmetric cleavage at the first division. All 19 tested embryos reached the blastocyst stage (Fig. 3D). Live-cell imaging revealed that Doubled-Dip exhibited cell cycle and mitotic abnormalities similar to those in Hap. They entered the second mitosis after several hours of delay and showed prolonged mitosis compared to Doubled-Tet (Fig. S3E-F). Moreover, Doubled-Dip had a higher incidence of abnormalities during the second and third mitoses (Fig. 3E-F; Video S3), mainly cytokinesis failure leading to binucleated blastomeres. As in Hap, 40.0% of Doubled-Dip blastomeres with second mitosis defects arrested before the third division, and 20.0% exhibited additional abnormalities during the third. Among those completing the second mitosis, 44.0% showed defects in the third. Tetraploidization caused by cytokinesis failure, similar to diploidization in Hap, did not improve development. Together, these findings indicate that halving the D/C ratio significantly impairs preimplantation development, even in diploid embryos. Embryos with a relative D/C ratio of 0.5 consistently showed a prolonged 2-cell stage and frequent mitotic defects, primarily cytokinesis failure starting at the second cleavage.

### Developmental defects also occur when the D/C ratio is halved at the late G2 phase of the 1-cell stage

In previous experiments, we manipulated the D/C ratio at the onset of development. To determine whether a proper D/C ratio is required throughout preimplantation development or only during specific stages, we next reduced the D/C ratio at later time points. During cleavage, embryonic cells divide without growing, so the D/C ratio is lowest at the 1-cell stage and doubles with each division. A relative D/C ratio of 0.5 at the 1-cell stage is thus not normally encountered and may impair progression. To examine whether the unusually low D/C ratio at the 1-cell stage impairs development, Dip was cultured to the late G2 phase of the first cell cycle, and one of the two pronuclei was removed, generating haploid embryos (hereafter 1G2-D/C^0.5^) (Fig. 4A). To ensure enucleation just before the first cleavage, which begins around 14 hpa, mitotic entry was blocked with a CDK1 inhibitor, and enucleation was performed between 13 and 15 hpa. Upon inhibitor release, over 90% of the embryos entered mitosis and reached the 2-cell stage without detectable mitotic defects (Fig. 4B-C, S4A; Video S4). Sham control embryos, subjected to identical treatment and microneedle insertion but without enucleation, developed efficiently to the blastocyst stage. In contrast, 1G2-D/C^0.5^ exhibited a trajectory resembling that of Hap. By Day 2, approximately 45% arrested at the 2–3-cell stage, and only 18.4% formed blastocysts (Fig. 4C-D). Live-cell imaging further revealed delayed entry into the second mitosis, prolonged mitotic progression (Fig. S4B-C), and mitotic abnormalities similar to those seen in Hap and Doubled-Dip (Fig. 4E-F; Video S5).

**Figure 4.**
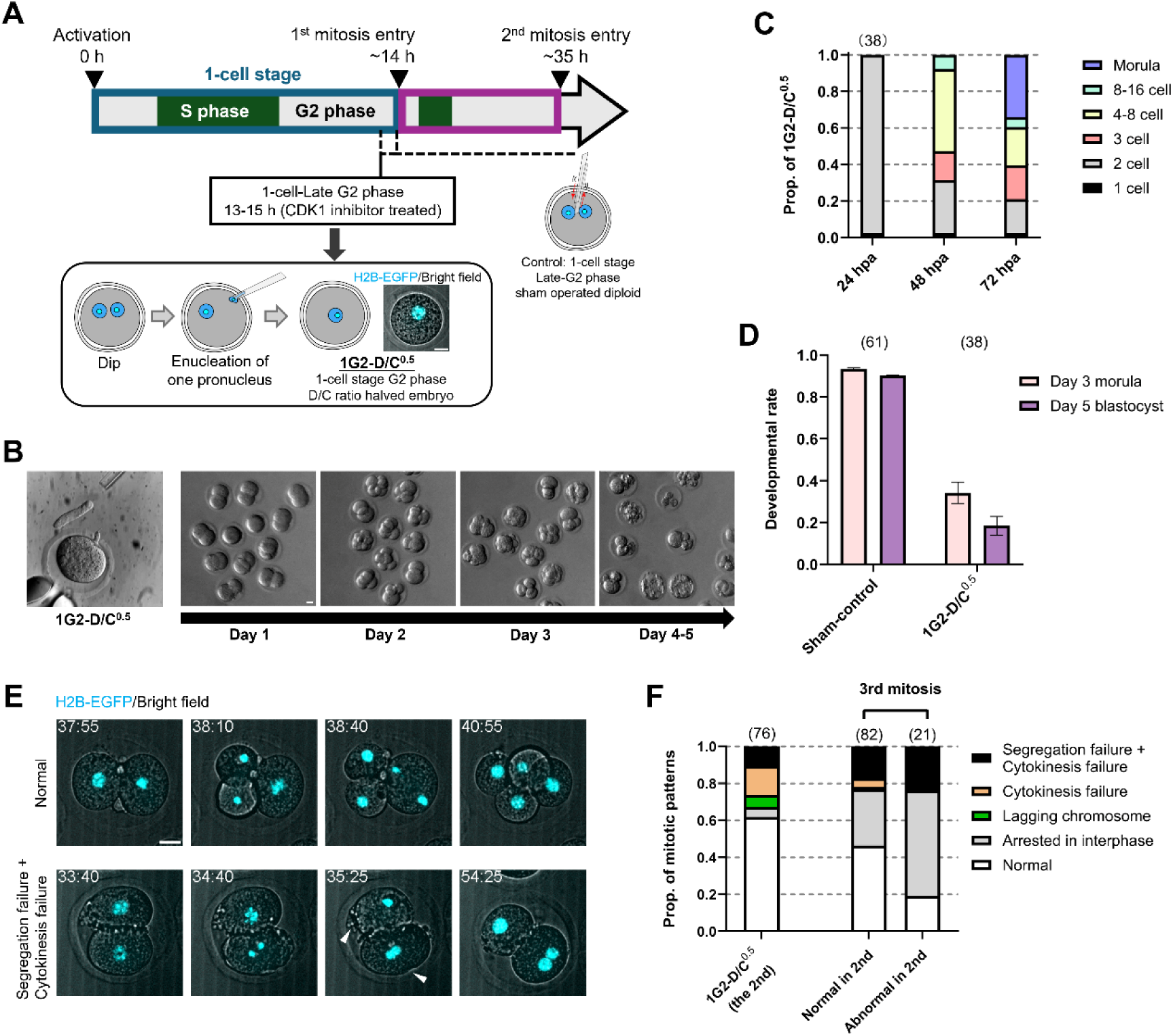
Postponing D/C ratio halving from the second meiosis to late G2 phase in 1-cell stage fails to rescue development. (A) Schematic of experimental design. Bottom right: Representative still image from time-lapse imaging of 1-cell stage G2 phase D/C ratio halved embryo (1G2-D/C^0.5^). (B) Bright-field images of 1G2-D/C^0.5^ after enucleation (left) and during culture. (C) Embryo statuses at 24, 48, and 72 hpa (≥4 experiments). (D) Proportions of embryos reaching the morula (Day 3) and blastocyst (Day 5) stages (mean ± SEM; ≥4 experiments). (E) Still images from time-lapse imaging of the second mitosis in 1G2-D/C^0.5^ expressing histone-H2B-EGFP (cyan, z-projected). Timestamps indicate hpa. White arrows mark incomplete cleavage furrows. See also Video S5. (F) Second mitosis outcomes in blastomeres from 1G2-D/C^0.5^ (left) and third mitosis outcomes categorized by second mitosis status (right) (≥3 experiments). Scale bars, 20 μm. Sample sizes are shown in parentheses above each bar.

Notably, nuclei in 2-cell-stage blastomeres of 1G2-D/C^0.5^ were smaller than those in Dip or Hap embryos (Fig. S4D, E). Although nuclear size did not differ significantly between blastomeres with or without mitotic abnormalities in the subsequent division (Fig. S4F), reduced nuclear size may still have contributed to impaired mitosis and lower developmental potential.

### Halving the D/C ratio at the late 2-cell stage does not impair preimplantation development

To evaluate the effects of reducing the D/C ratio at later developmental stages without altering nuclear size, we halved the D/C ratio at several time points during the 2-cell stage. This was achieved by enucleating one blastomere and fusing it with the other to form a single cell (Fig. 5A). Cell cycle progression in 2-cell-stage Dip embryos was determined by BrdU incorporation (Fig. S4G–H). Based on this, the D/C ratio was reduced during early S phase (17–19 hpa), early G2 phase (22–29 hpa), and late G2 phase (35–36 hpa, following CDK1 inhibition), generating 2S-D/C^0.5^, 2EarlyG2-D/C^0.5^, and 2LateG2-D/C^0.5^, respectively (Fig. 5A). To account for possible procedural effects, three control groups with a relative D/C ratio of 1.0 were designed and established. These included: (Control 1) electrofusion of two blastomeres to generate tetraploid embryos at the early S phase; (Control 2) removal of one blastomere followed by electrical stimulation of the other; and (Control 3) 2-cell-stage Dip treated with CDK1 inhibitor in the late G2 phase (Fig. 5A). All control embryos efficiently developed to the blastocyst stage (Fig. 5B-D), indicating that the procedures themselves did not significantly affect development. In contrast, 2S- D/C^0.5^ frequently arrested before the morula stage, with 52.6% reaching the morula and 36.8% the blastocyst stage. These rates were modestly higher than those of Hap and Doubled-Dip, but still significantly lower than embryos with a relative D/C ratio of 1.0 (Fig. 5B–D). Moreover, live-cell imaging revealed that approximately 20% of 2S-D/C^0.5^ failed to complete cytokinesis during the second mitosis (Fig. 5E-F; Video S6), resembling defects observed in other halved D/C ratio embryos such as Hap and Doubled- Dip. These findings indicate that even when the D/C ratio is normal during the 1-cell stage, its reduction from the early 2-cell stage can induce defects starting at the second mitosis, ultimately leading to developmental arrest. Notably, developmental efficiency improved as the timing of D/C ratio halving was delayed within the 2-cell stage, reaching control levels in 2LateG2-D/C^0.5^ (Fig. 5D). This suggests that maintaining a normal D/C ratio until the late G2 phase of the 2-cell stage is sufficient to support normal preimplantation development.

**Figure 5.**
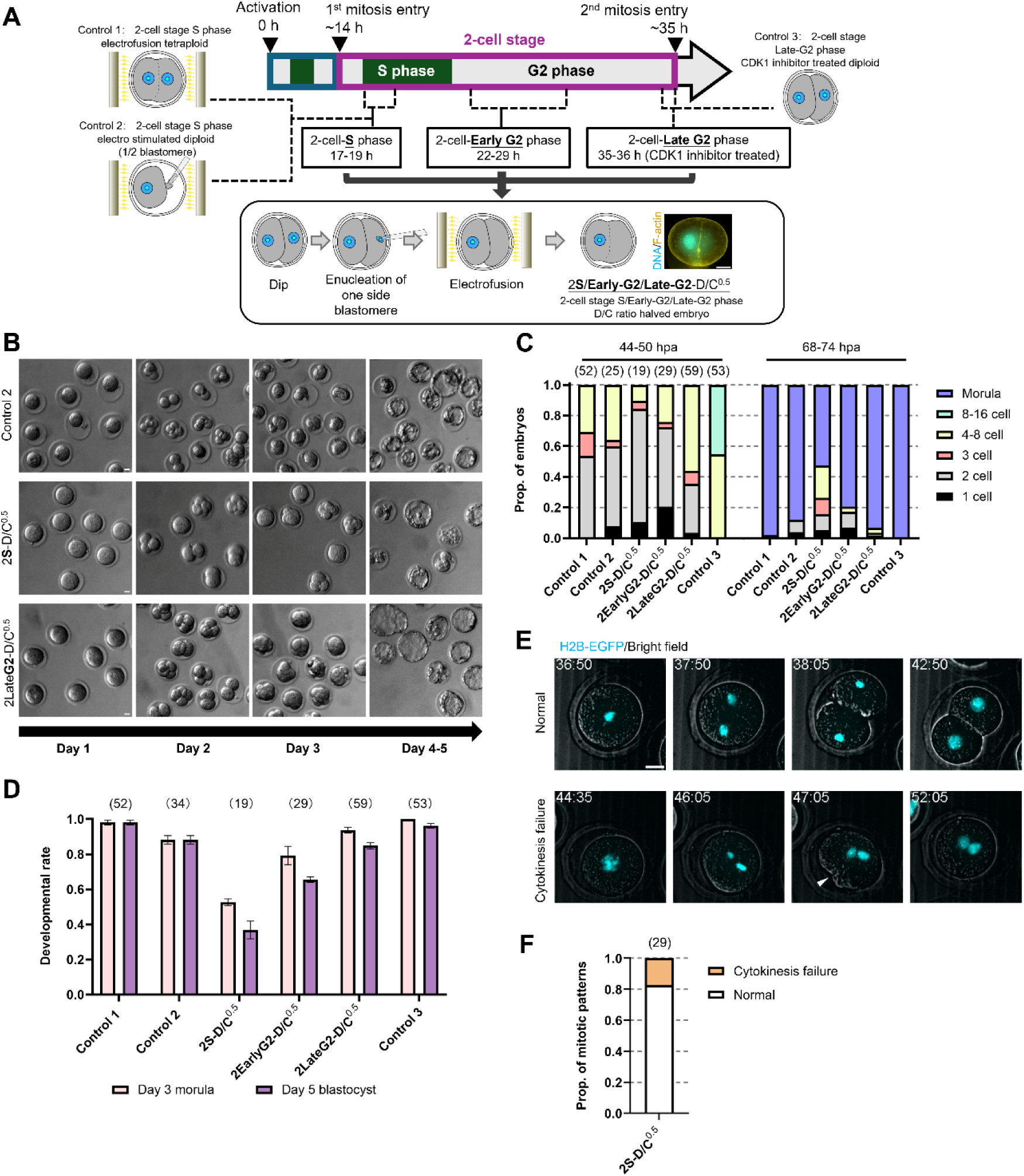
Postponing D/C ratio halving from the 1-cell stage to the 2-cell stage alleviates developmental defects. (A) Schematic of experimental design. Bottom right: representative immunofluorescence image of 2-cell stage S phase D/C ratio halved embryo (2S-D/C^0.5^). (B) Bright-field images of Control 2 embryos (as described in A, top), 2S-D/C^0.5^ and 2- cell stage late G2 phase D/C ratio halved embryo (2LateG2-D/C^0.5^). (C) Embryo statuses at 24, 48, and 72 hpa (≥3 experiments). (D) Proportions of embryos reaching morula (Day 3) and blastocyst (Day 5) stages (mean ± SEM; ≥3 experiments). (E) Still images from time-lapse imaging of the second mitosis in 2S-D/C^0.5^ expressing histone-H2B-EGFP (cyan, z-projected). Timestamps indicate hpa. White arrows mark incomplete cleavage furrows. See also Video S6. (F) Second mitosis outcomes in blastomeres from 2S-D/C^0.5^ based on three independent experiments. Scale bars, 20 μm. Sample sizes are shown in parentheses above each bar.

Taken together, our findings demonstrate that the DNA-to-cytoplasm (D/C) ratio is critical for mouse preimplantation development, particularly during the 2-cell stage. Halving the D/C ratio at this stage results in mitotic abnormalities, such as spindle defects and cytokinesis failure, and is accompanied by increased developmental arrest.

## Discussion

In placental mammals, both biparental genomic contribution and diploidy are essential for successful embryonic development. Embryos with a diploid genome of uniparental origin, or a biparental genome in a non-diploid state, fail to support post-implantation development^30–34^. In contrast, these constraints do not apply during preimplantation development. Diploid embryos with only maternal or paternal genomes, as well as polyploid embryos, can develop efficiently to the blastocyst stage^5,35^. However, haploid embryos, whether of maternal or paternal origin, exhibit markedly low developmental rates, with most arresting before the blastocyst stage for reasons that remain unclear. In this study, by generating embryos with varying cell sizes and genome ploidies, we show that while haploidy may contribute to developmental failure, the primary cause of arrest is an imbalance between genomic DNA content and cytoplasmic volume.

Cytoplasmic aspiration with a glass capillary is a standard method to reduce the size of mouse oocytes and early embryos^15,28,36,37^. Here, we instead used a non-invasive approach that alters spindle positioning to convert the second meiotic division from asymmetric to symmetric, thereby reducing cytoplasmic volume in one-cell embryos. We analyzed embryos that divided nearly equally, yielding cells with about half the original volume (Fig. S1A). As this method also produces embryos with varying asymmetry, it can generate a range of D/C ratios.

The higher developmental rates of Halved-Hap and Halved-Hap-1 compared to Hap suggest that although diploidy and oocyte size are considered important for preimplantation development, maintaining the balance between genomic content and cytoplasmic volume can alleviate developmental impairment. Nevertheless, their developmental rates remain lower than that of Dip, indicating that haploidy may impose additional limitations even when the D/C ratio is normalized. The higher blastocyst formation rate in Halved-Hap compared to Halved-Hap-1 implies that starting development with a greater number of blastomeres may help offset reduced proliferation associated with haploidy. Moreover, despite diploidy, Doubled-Dip exhibited low blastocyst formation rates. These rates were restored by inducing tetraploidy, indicating that simply increasing cytoplasmic volume does not improve preimplantation development. Rather, this highlights the importance of maintaining a proper balance between genome content and cytoplasmic volume or cell size. Similar findings have been reported in studies using embryos produced by fusing multiple oocytes, where increased cytoplasmic volume was associated with reduced developmental success^16,38^.

The primary developmental barrier in Hap arises before the morula stage, beginning at the 2-cell stage. Although over 30% of Hap appear morphologically arrested at the 2-cell stage (Fig. 2A), live imaging showed that nearly all blastomeres had entered the second mitosis, despite a delayed interphase (Fig. 2F). However, many failed to complete cytokinesis, with or without preceding chromosome segregation failure, and thus retained a 2-cell-like appearance. Cytokinesis failure has also been reported in haploid somatic cells, often resulting in diploidization. Notably, diploidized somatic cells exhibit greater stability and proliferative capacity than haploid cells^39,40^. In contrast, most blastomeres in Hap embryos undergoing cytokinesis failure either arrested in the next cell cycle or accumulated further mitotic errors (Fig. 2F). These differences suggest that the mechanisms of cytokinesis failure differ between Hap blastomeres and haploid somatic cells, which appear to maintain a D/C ratio similar to diploid cells^41^. In Hap blastomeres, the defects are more likely caused by a reduced D/C ratio, as similar abnormalities were observed in Doubled-Dip.

What causes the mitotic and cytokinesis abnormalities in embryos with a reduced D/C ratio remains unclear. One possibility is that chromosome mis-segregation arises from DNA damage–induced fragmentation caused by a shortened G2 phase at the 2-cell stage^42^. However, in our study, the 2-cell stage was prolonged (Fig. S2A), and DNA damage markers were not significantly elevated in halved-D/C embryos (Fig. S2C, D), suggesting that DNA damage is unlikely to be the primary cause.

Although the precise mechanism remains to be defined, our findings collectively suggest how a reduced D/C ratio may compromise mitotic fidelity. Notably, embryos with a halved D/C ratio showed no mitotic abnormalities during the first division (Fig. 2D, Fig. S4A), when the cell is at its largest. This implies that mitotic errors are not due to imbalance between DNA content and cell size *per se*, but rather stem from insufficient concentrations of key cytoplasmic factors. Moreover, Cytokinesis failure occurred in only 20–30% of blastomeres and, even in these cases, a cleavage furrow formed transiently but failed to complete (Fig. 2E, 3E; Video S1, S3). Similarly, chromosome segregation failure, observed in 10–30% of blastomeres, was consistently associated with malformed spindles (Fig. 2G-I, Fig. S2E-G) and cytokinesis failure. These data suggest that the spindle and cytokinesis machinery are largely intact but function suboptimally due to dilution of limiting factors. Our findings also indicate that an appropriate D/C ratio during the 2-cell stage—when major ZGA begins and maternal factors decline—is essential for accurate mitosis and continued development.

Previous studies in budding yeast and human cell lines have demonstrated that an abnormally low D/C ratio causes genome dilution, which, in severe cases, leads to a collapse in biosynthetic activity ^18,19^. A recent report in *C. elegans* further validated that although transcription is partially upregulated under low D/C conditions, intracellular mRNA concentrations remain diluted, with highly expressed transcripts being particularly vulnerable to this effect^43^. Building on this evidence, we speculate that in 2-cell-stage mouse embryos, certain proteins that require a minimum cytoplasmic concentration, such as those involved in spindle assembly and cytokinesis, may become limiting when the D/C ratio is halved. These limitations may compromise mitotic fidelity and, alongside other cytoplasmic insufficiencies, may underlie the observed developmental arrest. This model accounts for the absence of defects during the first mitosis, when maternal factors are still abundant, and explains why reducing the D/C ratio later in the 2-cell stage improves development by providing more time for the synthesis of essential embryonic factors.

Binucleated blastomeres with equally sized nuclei are relatively common in human embryos generated by *in vitro* fertilization^44–48^. Some of these are believed to result from cytokinesis failure, although the underlying mechanism remains unclear. A previous study reported that, in humans—where ZGA occurs at the 8-cell stage—approximately 65% of embryos at the 9- to 16-cell stage contain one to six binucleated blastomeres. A subset of these blastomeres subsequently arrest during later cleavage divisions^49^. Such post-ZGA binucleation may involve mechanisms similar to those underlying mitotic abnormalities observed in our study.

Compared to other vertebrates, mammalian oocytes have a relatively small volume^9^. However, because ZGA occurs after fewer cell divisions, blastomeres remain considerably large at the time of ZGA^22^. Under these conditions, excess cytoplasm may dilute the concentration of certain embryonic factors, causing blastomeres to enter mitosis before they are fully prepared. This can result in mitotic errors and impaired development. Further investigation of this mechanism will enhance our understanding of how the balance between genomic content and cytoplasmic volume is regulated during preimplantation development in mammals.

## Resource Availability

This study did not generate any unique datasets or code.

Any additional information required to reanalyze the data reported in this paper is available from the lead contact upon request.

## Acknowledgments

We thank Tomo Kondo and Ryohei Nakamura for valuable discussions and insightful comments on the manuscript. We are grateful to Hiroshi Kimura for providing the plasmid encoding mRFP1-tagged histone H2B, and to Kazuo Yamagata for the pcDNA3.1-pA vector. We also thank Hirohisa Kyogoku and Hiroyuki Imai for their technical advice on enucleation and electrofusion of mouse embryos.

This work was supported by the Japan Society for the Promotion of Science (JSPS) KAKENHI (Grants 18K19325, 20H05356, 22H04664, and 23H02485). The first author, T.P., was supported in 2024 by JST SPRING (Grant Number JPMJSP2108).

## Author Contributions

T.P. and M.O. conceived and designed the study. N.T. and T.P. developed the method to generate half-sized embryos, performed immunofluorescence and live imaging of haploid embryos, and analyzed the corresponding data. T.P. conducted all other experiments and performed the remaining data analyses. T.P. drafted the manuscript. M.O. supervised the project, reviewed and revised the manuscript, and secured funding.

## Declaration of Interests

The authors declare no competing interests.

### Declaration of Generative AI and AI-assisted Technologies in the Writing Process

The author used ChatGPT (OpenAI, GPT-4 Turbo) to assist with language refinement, including improvements to clarity, grammar, and readability. All content was critically reviewed and approved by the author, who assumes full responsibility for the final manuscript.

## Key resources table

| REAGENT or RESOURCE | SOURCE | IDENTIFIER |
| --- | --- | --- |
| Antibodies |  |  |
| Alpha Tubulin (YOL1/34), Rat mAb | Santa Cruz<br>Biotechnology | sc-53030; RRID:<br>AB_2272440 |
| Lamin B1 (B-10) Alexa Fluor® 488, Mouse mAb | Santa Cruz<br>Biotechnology | sc-374015; RRID:<br>AB_10947408 |
| phospho-Histone H2A.X (Ser139), Mouse mAb | Merck Millipore | JBW301; RRID:<br>AB_309864 |
| RPA32/RPA2 (4E4), Rat mAb | Cell Signaling<br>Technology | 2208T; RRID:<br>AB_2238543 |
| BrdU, Rat mAb | BioRad | OBT0030; RRID:<br>AB_2314029 |
| Goat anti-Mouse IgG (H+L) Highly Cross-Adsorbed<br>Secondary Antibody, Alexa Fluor™ 488 | Invitrogen | A-11029; RRID:<br>AB_2534088 |
| Goat anti-Mouse IgG (H+L) Highly Cross-Adsorbed<br>Secondary Antibody, Alexa Fluor™ 555 | Invitrogen | A-21424; RRID:<br>AB_141780 |
| Goat anti-Mouse IgG (H+L) Highly Cross-Adsorbed<br>Secondary Antibody, Alexa Fluor™ 647 | Invitrogen | A-21236; RRID:<br>AB_2535805 |
| Donkey anti-Rat IgG (H+L) Highly Cross-Adsorbed<br>Secondary Antibody, Alexa Fluor™ 488 | Invitrogen | A-21208; RRID:<br>AB_141709 |
| Donkey anti-Rat IgG (H+L) Highly Cross-Adsorbed<br>Secondary Antibody, Alexa Fluor™ 568 | Invitrogen | A78946; RRID:<br>AB_2910653 |
| Donkey anti-Rat IgG (H+L) Highly Cross-Adsorbed<br>Secondary Antibody, Alexa Fluor™ 647 | Invitrogen | A78947; RRID:<br>AB_2910635 |
| Chemicals, peptides, and recombinant proteins |  |  |
| Rhodamine phalloidin | Invitrogen | R415; |
| Hoechst 33342 dye | Thermo Fisher<br>Scientific | H-1399 |
| Pregnant mare serum gonadotropin (PMSG) | ZENOAQ | N/A |
| Human chorionic gonadotropin (hCG) | Kyoritsu Seiyaku | N/A |
| Hyaluronidase from bovine testes | Sigma-Aldrich | H4272 |
| M16 medium | Sigma-Aldrich | Cat# M7292 |
| M2 medium | Sigma-Aldrich | Cat# M7167 |
| Tyrode's solution | Sigma-Aldrich | Cat# T1788 |
| Cytochalasin B | Sigma-Aldrich | Cat# C6762 |
| DMSO | WAKO | Cat# 048-21985 |
| Ribo m<sup>7</sup>G Cap Analog | Promega | Cat# P1711 |
| Paraffin oil | Nacalai Tesque | Cat# 26137-85 |
| Ro-3306 | COSMO BIO | Cat# HY-12529 |
| CK-666 | Selleck | Cat# S7690 |
| Phytohemagglutinin-P | WAKO | Cat# 161-15251 |
| Critical commercial assays |  |  |
| RiboMax<sup>TM</sup> Large Scale RNA Production System-T7 | Promega | Cat# P1320 |
| Experimental models: Organisms/strains |  |  |
| Mouse/CB6F1(BALB/cA x C57B/6J) |  | N/A |
| Mouse/BALB/cA (wild type) | CLEA Japan | https://www.clea-japan.com/products/inbred/item_a0430 |
| Mouse/C57B/6J (wild type) | CLEA Japan | https://www.clea-japan.com/products/inbred/item_a0420 |
| Mouse/R26-H2B-EGFP (Accession No.CDB0238K) | RIKEN Center for Biosystems Dynamics Research | CDB0238K |
| Mouse/BALB/cA-R26-H2B-EGFP N4+ | This paper | N/A |
| Mouse/C57BL/6J-R26-H2B-EGFP N4+ | This paper | N/A |
| Recombinant DNA |  |  |
| Plasmid: pcDNA3.1-pA for mRNA production | Dr. Kazuo Yamagata | N/A |
| Plasmid: mRFP1-tagged histone H2B | Dr. Hiroshi Kimura | N/A |
| Plasmid: EGFP-tagged α-tubulin | This paper | N/A |
| Software and algorithms |  |  |
| Fiji (ImageJ) | National Institutes of Health | https://imagej.nih.gov/ij/ |
| Microsoft Excel | UTokyo Microsoft License | https://utelecon.adm.u-tokyo.ac.jp/microsoft/ |
| GraphPad Prism 10 | GraphPad Software | https://www.graphpad.com/ |
| Adobe Illustrator | Adobe Creative Cloud | https://www.adobe.com/jp/creativecloud.html |
| Other |  |  |
| Delta Vision Core System | Applied Precision | N/A |
| Olympus IX71 microscope (20x/0.85 NA oil objective lens) | Olympus | N/A |
| Olympus IX70 microscope (20x/0.85 NA oil objective lens) | Olympus | N/A |
| Olympus IX70 microscope (60x/1.30 Sil oil objective lens) | Olympus | N/A |
| CSU10 | Yokogawa | N/A |
| CSU-X1 | Yokogawa | N/A |
| CellVoyager CV1000 | Yokogawa | N/A |
| Piezo-driven micromanipulator | Prime Tech | N/A |
| LF101 electric cell fuser | BEX Co. | N/A |
| 1-mm fusion chamber | BEX Co. | N/A |
| P-1000IVF | Sutter Instrument | N/A |
| Microforge MF-900 | Narishige | N/A |

## Methods

### Mice

Unless otherwise noted, CB6F1 mice, generated by crossing female BALB/cA and male C57BL/6J mice (CLEA Japan), were used for oocyte collection. R26-H2B-EGFP mice (Accession No. CDB0238K), provided by the RIKEN Center for Biosystems Dynamics Research^50^, were backcrossed with BALB/cA and C57BL/6J mice for more than four generations to generate R26-H2B-EGFP mice with a CB6F1 background. Oocytes from R26-H2B-EGFP mice with a CB6F1 background were also used in some experiments. All mice were group-housed under standard conditions with a 12-hour light/dark cycle and had ad libitum access to food and water. All animal experiments were approved by the Animal Experimentation Committees of the Graduate School of Arts and Sciences (approval no. 26-29) and the Graduate School of Science (approval no. A2024S010), The University of Tokyo, and conducted in accordance with the guidelines for animal use issued by the Committee of Animal Experiments at The University of Tokyo.

### Superovulation and collection of metaphase II oocytes

Female CB6F1 mice (8–16 weeks old) were superovulated by intraperitoneal injection of 5 IU PMSG (ZENOAQ), followed 48 hours later by 5 IU hCG (Kyoritsu Seiyaku). Cumulus–oocyte complexes were collected from the oviduct 16–20 hours after hCG injection. Cumulus cells were removed using M2 medium (Sigma-Aldrich) containing 100 μg/mL hyaluronidase. Denuded oocytes were washed and cultured in M16 medium (Sigma-Aldrich) at 37°C in 5% CO₂. Unless otherwise stated, media were prepared as microdrops under paraffin oil (Nacalai Tesque) in Petri dishes. Oocytes showing fragmentation, spontaneous activation, atrophy, or large perivitelline space were excluded.

### Sr²⁺-induced parthenogenetic activation

Sr²⁺-induced parthenogenetic activation was performed using M16 medium supplemented with 5 mM SrCl₂ and 5 mM EGTA. For Dip embryo generation, 2.5 μg/mL cytochalasin B (Sigma-Aldrich) was added to inhibit second polar body extrusion. For Hap embryos, Metaphase-II oocytes were preincubated in M16 with 250 μM CK-666 (Selleck) for 3 hours, then transferred to activation medium containing the same concentration^51^. To prevent CK-666 diffusion into paraffin oil and ensure consistent concentration, all CK-666 treatments were performed in 96-well plates without oil.

### Electrofusion and electrical parthenogenetic activation

Fusion of Meta-II oocytes was performed following a protocol adapted from a previous study^37^. Oocytes were treated with acid Tyrode’s solution (Sigma-Aldrich) to remove the zona pellucida, followed by incubation in M2 medium containing 50 μg/mL phytohemagglutinin (WAKO) for 1 minute. Pairs of oocytes were then aligned under a micromanipulator using a blunt needle to ensure close apposition. The paired oocytes were transferred to a 1-mm fusion chamber (BEX Co.) connected to the LF101 electric cell fuser (BEX Co.). Fusion was induced using 5 V alternating current at 0.5 MHz for 5 seconds, followed by a 60 V direct current pulse for 50 μs. Using M2 medium in the chamber resulted in simultaneous fusion and parthenogenetic activation. After electrofusion, the oocytes were transferred to M16 medium with or without 2.5 μg/mL cytochalasin B, depending on the experimental condition, to generate Doubled-Tet or Doubled-Dip, respectively. For the fusion or stimulation of blastomeres in 2-cell-stage embryos, embryos were subjected to 5 V alternating current at 0.5 MHz for 5 seconds, followed by a 160 V direct current pulse for 40 μs in an M2 medium-filled fusion chamber. After electrofusion, the embryos were transferred to M16 medium for subsequent culture.

### Enucleation

Embryos were placed in M2 medium containing 5 μg/mL cytochalasin B for approximately 10 minutes prior to enucleation. A glass needle (5–10 μm in diameter) was used to penetrate the plasma membrane under a micromanipulator and Piezo Impact Drive (Prime Tech). The nucleus was aspirated and gently removed. To perform enucleation during the late G2 phase, embryos were transferred to M16 medium containing 20 μM CDK1 inhibitor Ro-3306 (COSMO BIO) two hours before the average time of mitotic entry, as determined by live-cell imaging in this study. After enucleation, embryos were transferred to M16 medium containing 20 μM Ro-3306 for at least 1 hour to allow recovery from manipulation, followed by washing in M16 medium to remove the inhibitor before further culture. Sham operations for late G2 phase enucleation were conducted under identical conditions, including drug treatment and room temperature exposure. For late G2 phase embryos at the 1-cell stage, sham operations involved piercing the region between the two pronuclei 2–3 times using needles of the same diameter as in the experimental group. For 2-cell stage late G2 phase embryos, sham operations followed the same inhibitor treatment and room temperature exposure as the experimental group.

### mRNA preparation and microinjection

mRNAs were synthesized *in vitro* from linearized template plasmids using the RiboMax™ Large Scale RNA Production System-T7 (Promega), supplemented with Ribo m⁷G Cap Analog (Promega). Reaction mixtures were treated with DNase I after RNA synthesis. Synthesized RNAs were purified by phenol extraction followed by ethanol precipitation^52^. The template plasmid for EGFP-tagged α-tubulin was constructed by inserting a DNA fragment encoding EGFP–α-tubulin into the pcDNA3.1-polyA vector^53^. Microinjection was performed with the following mRNAs: mRFP1-tagged histone-H2B at 50 ng/μL and EGFP-tagged α-tubulin at 200 ng/μL. Minimal volumes (picoliters) of mRNA were injected into Meta-II oocytes using a Piezo-driven micromanipulator (Prime Tech) with glass needles approximately 3 μm in diameter.

### Live imaging

Confocal imaging was performed using an Olympus IX71 microscope (20×/0.85 NA oil objective lens) equipped with a spinning disk confocal system (CSU10 or CSU-X1; Yokogawa) and a CO₂ microscope stage incubator (TOKAI HIT). Images were captured with a CCD camera (iXon DU897E-CSO-#BV; ANDOR) controlled by MetaMorph software (Universal Imaging). In imaging experiments involving EGFP–α-tubulin, some observations were conducted using the CellVoyager CV1000 confocal scanner box (Yokogawa). Images were captured with a CCD camera (iXon DU897E-CSO-#BV; ANDOR) managed by Metamorph software (Universal Imaging). In imaging experiments involving EGFP-α-tubulin, some were conducted using the confocal scanner box CellVoyager CV1000 (Yokogawa). Time-lapse imaging included a single mid-plane bright-field reference and one or two fluorescence channels acquired at 4-μm intervals (21 Z-sections). Images were shown as maximum intensity projections, with contrast-adjusted bright-field overlays. Time intervals ranged from 5 minutes (short time-lapse for the first mitosis) to 15 minutes (long time-lapse from the second mitosis onward). As 1- cell stage embryos were highly light-sensitive and often failed to develop beyond the 2- cell stage, imaging of the first and subsequent mitoses was performed separately.

### BrdU incorporation assay

Diploid embryos were transferred to M16 medium containing 0.2 mM BrdU at 4, 8, 12, 16, 18.5, 20, and 24 hpa. After 1 hour of incubation, embryos were fixed with 4% paraformaldehyde.

### Immunofluorescence

After removal of the zona pellucida using acidic Tyrode’s solution, oocytes or embryos were washed with 0.5% polyvinyl pyrrolidone in PBS and subsequently fixed in 4% paraformaldehyde for 30–40 minutes at 25°C. Fixed samples were permeabilized with 0.1% Triton X-100 in PBS for 10–15 minutes at 25°C, followed by blocking in PBS containing 3% BSA and 0.1% Tween-20 for 1 hour at 25°C or overnight at 4°C. For BrdU staining, cells were permeabilized with 2 M HCl and 0.5% Triton X-100 in PBS for 45 minutes. After blocking, samples were sequentially incubated with primary antibody solution for 1 hour at 25°C or overnight at 4°C, followed by incubation with secondary antibody solution containing Hoechst 33342 (Thermo Fisher Scientific) and rhodamine-phalloidin (Invitrogen) for 1 hour at 25°C. Each incubation step was followed by three 5- minute washes in PBS containing 0.1% BSA and 0.1% Tween-20. Samples were observed at 25°C using a fluorescence microscope (IX70, Olympus) equipped with 60×/1.35 NA or 20×/0.85 NA oil immersion objective lenses and a CoolSNAP HQ camera (Roper Scientific). The system was controlled by DeltaVision SoftWorx software (Applied Precision). Fluorescence images were acquired as Z-stacks at 1–2 μm intervals and displayed as maximum intensity Z-projections. The imaging medium consisted of PBS supplemented with 0.1% BSA.

### Image analysis

Image analysis was performed using ImageJ/Fiji. The volumes of blastomeres and nuclei were quantified by manually selecting their largest cross-sectional areas, estimating the area, and approximating the structures as spheres. The timing of nuclear envelope breakdown (NEBD) and anaphase onset, as well as the presence of chromosome segregation errors, were determined from fluorescence images of histone-H2B. The success of cytokinesis was assessed using bright-field images. Blastomeres that did not enter mitosis were excluded from cell cycle-related analyses. For blastomeres exhibiting severe chromosome segregation failure, anaphase onset was defined as the first frame showing evident chromosome displacement.

In live imaging, bright-field images were captured from a single slice near the middle of each sample and are presented as references to facilitate visualization of blastomere boundaries. Unless otherwise specified, fluorescence images represent Z-projections of the entire sample. For merged still images from live-cell imaging, the contrast of the bright-field channel was substantially reduced to better visualize the fluorescent signals. Unprocessed data can be found in the corresponding videos.

In the presentation of BrdU immunofluorescence images, the Z-projected BrdU signal was uniformly adjusted with a fixed contrast range (minimum: 500, maximum: 1500) to enhance the visibility of nuclear foci.

### Statistical analysis

Statistical analyses and graph preparation were performed using Microsoft Excel and GraphPad Prism 10. Sample sizes, statistical tests, and significance levels are indicated in the figures and figure legends.

---

**Figure S1.**
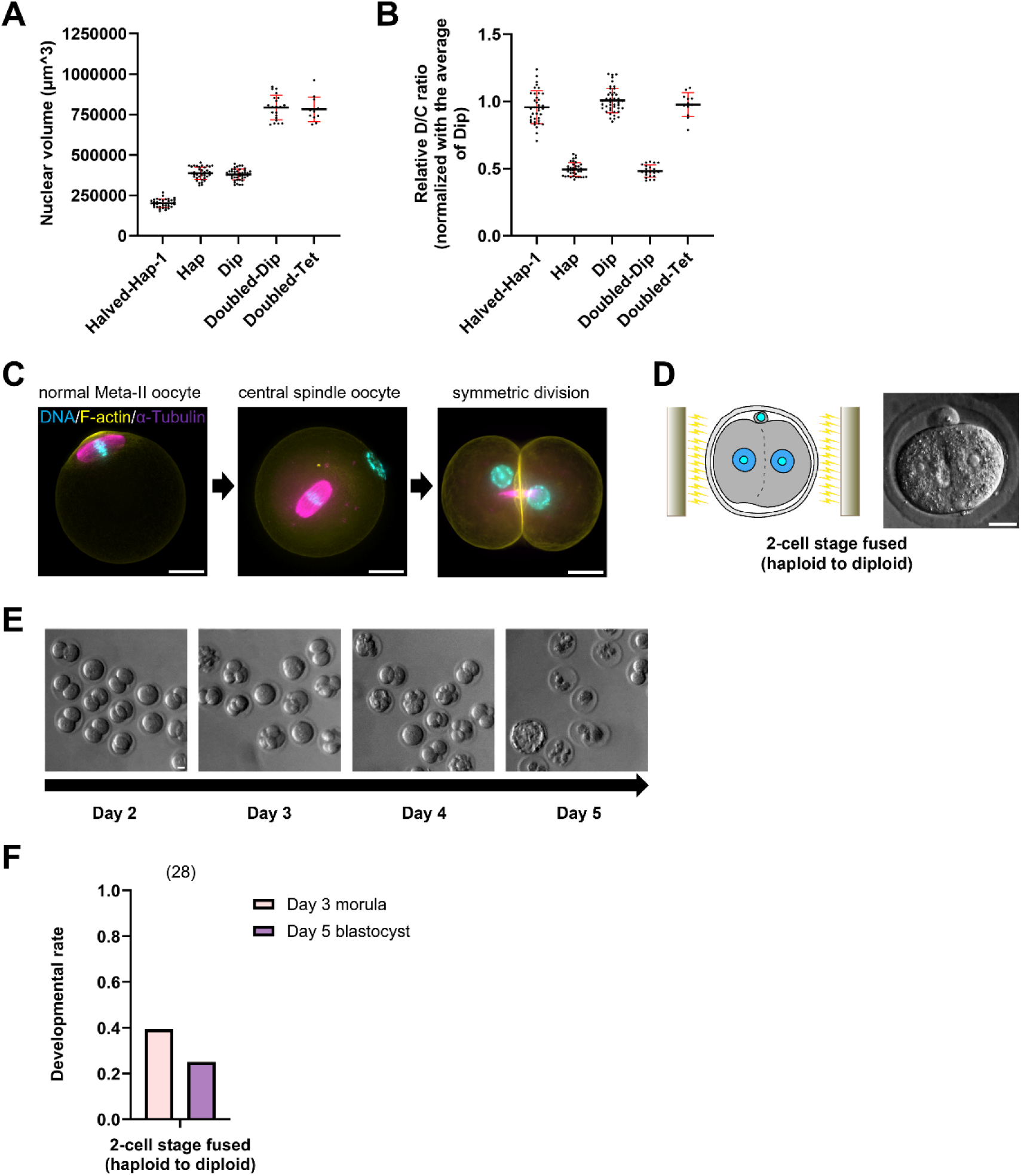
Validation of D/C ratio manipulation and evaluation of developmental outcomes in 2-cell-stage fused haploid embryos. Relates to Figure 1. (A and B) The volume of the embryos was calculated by manually outlining 1-cell stage embryos that appeared approximately circular and assuming them to be spheres. The D/C ratio was calculated by setting the ratio of the average volume of Dip to its ploidy as 1.0 (mean ± SD, number of embryos: n ≥ 12 from more than three independent experiments group). (C) Representative z-projected immunofluorescence images of Halved-Hap production. DNA is shown as cyan, LaminB1 is shown as red, and F-actin is shown as yellow. For the central spindle oocyte (middle), an image stack was acquired at approximately 30 μm from the middle part of the sample. Scale bars = 20 μm. (D and E) Schematic representation and representative bright-field images of 2-cell stage electrofused diploid embryo. Embryos were photographed approximately every 24 hours. (F) Day 3 morula rate and Day 5 blastocyst rate of 2-cell stage S phase electrofused diploid embryo (mean, samples from two independent experiments).

**Figure S2.**
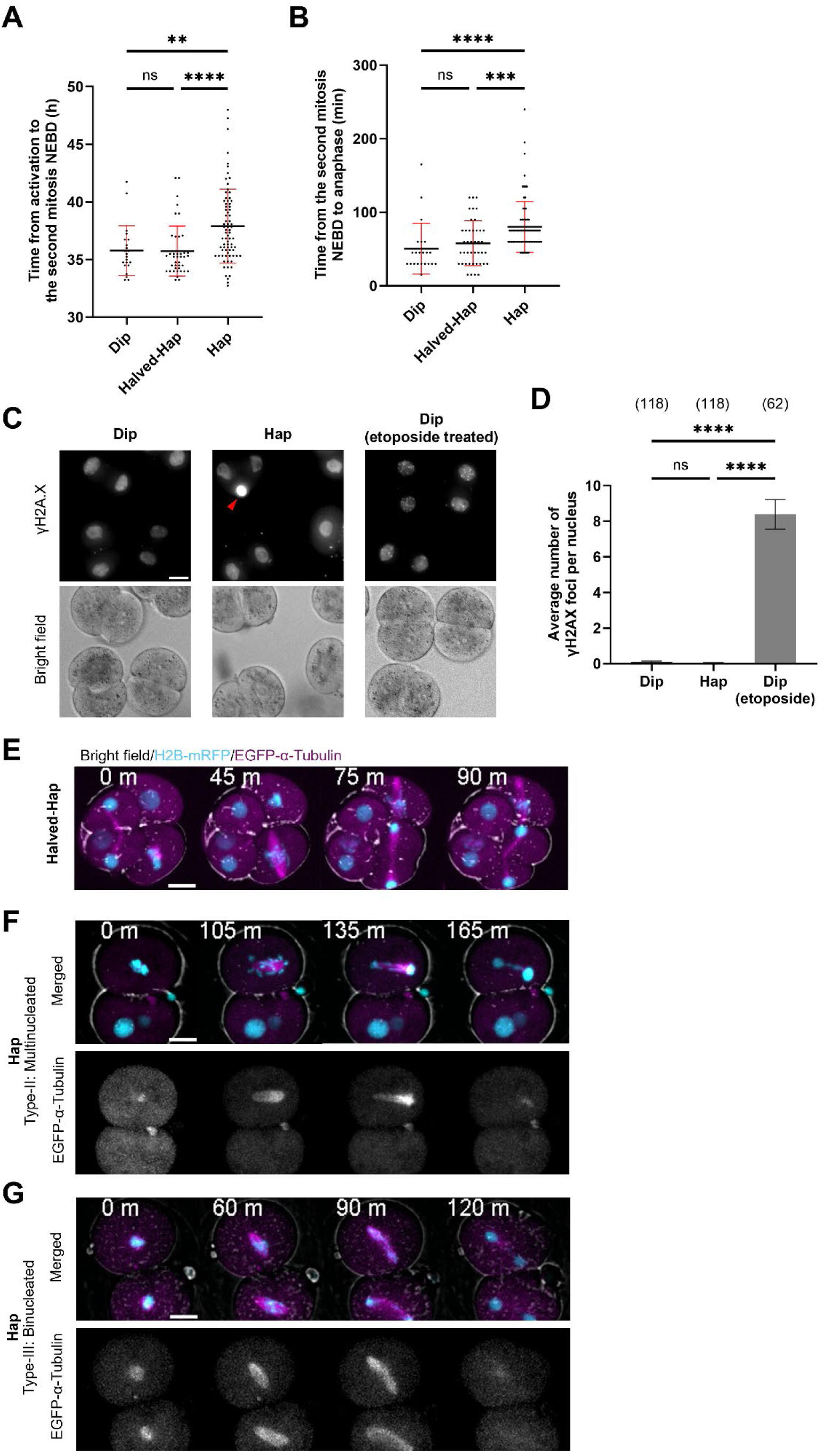
Haploid embryos exhibit delayed mitotic entry and spindle-associated abnormalities during the second mitosis without detectable DNA damage. Relates to Figure 2. (A and B) Scatterplots showing the time from activation to NEBD of the second mitosis, and the time from NEBD to anaphase during the second mitosis (mean ± SD, number of embryos: n ≥ 22 from more than three independent experiments group). Statistical analysis was performed using the Kruskal-Wallis test to confirm overall significant differences among groups, followed by pairwise comparisons using Dunn’s multiple comparisons test to account for multiple comparisons. (C) Representative immunofluorescence images showing z-projected γH2AX signals in Dip, Hap, and etoposide-treated Dip. For the etoposide-treated Dip group, embryos were exposed to 5 μM etoposide from 24 to 25 hpa. All samples were fixed at 30 hpa. Red arrow indicates the second polar body in Hap. (D) A quantified graph (mean ± SEM, samples from two independent experiments) showing the number of γH2AX foci in the samples mentioned in C. (E) Representative still images from time-lapse imaging of the second mitosis in Halved-Hap embryos. The embryos were microinjected with mRNAs encoding mRFP1-tagged histone-H2B (cyan, z-projected) and EGFP-tagged α-tubulin (magenta, z-projected) prior to parthenogenetic activation. See also Video S2. (F and G) Representative still images from time-lapse imaging of the second mitosis in Hap with cytokinesis failure Type-II (Multinucleated) and Type-III (Binucleated). See also related Video S2. It should be noted that the time-lapse imaging of Type-II and Type-III, as well as Halved-Hap, was captured using the confocal scanner unit CSU10, which has a slightly different resolution compared to the other time-lapse imaging data obtained with the confocal scanner unit CSU-X1 in this study. Scale bars = 20 μm, timestamps indicate the time from NEBD (in minutes) of the second mitosis, sample sizes are indicated in parentheses above each bar.

**Figure S3.**
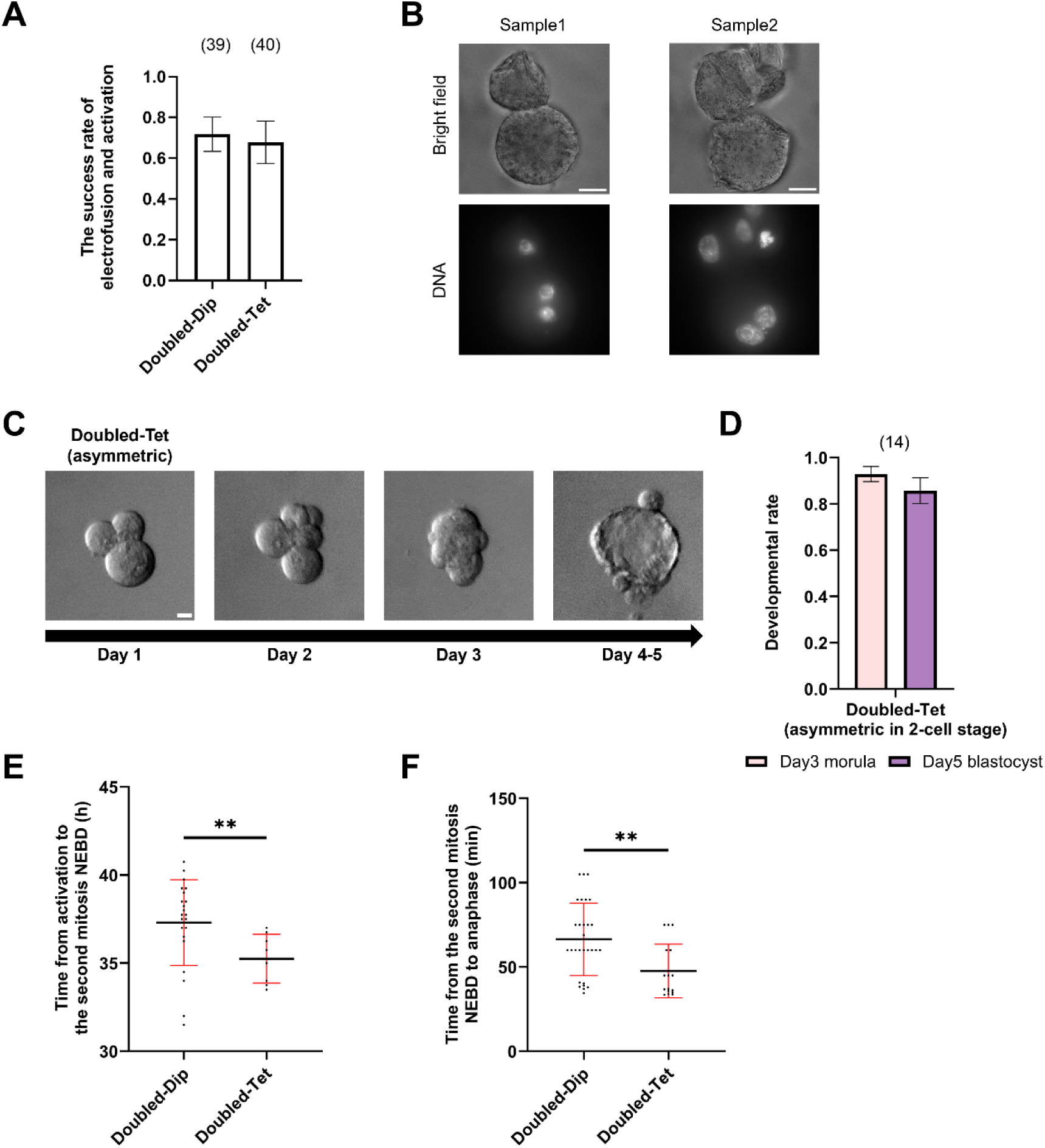
Electrofused diploid embryos exhibit abnormal nuclear morphology and delayed mitosis compared to tetraploid counterparts. Relates to Figure 3. (A) The success rate of electrofused and electro-activated embryos (mean ± SEM, samples from more than six independent experiments for each group). (B) Two fixed samples from arrested Doubled-Dip. DNA was stained with Hoechst and shown as a z-projected image. Since Doubled-Dip samples are scarce, only those that were confirmed to be unable to develop beyond the morula stage were selected for fixation. As a result, there may be a 48-hour interval between the occurrence of developmental arrest and fixation. Scale bars = 20 μm. (C) Representative bright-field images of Doubled-Tet with asymmetric cleavage on Day 1. Embryos were photographed approximately every 24 hours. (D) Day 3 morula rate and Day 5 blastocyst rate of asymmetric Doubled-Tet (mean ± SEM, samples from five independent experiments). (E and F) Scatterplots showing the time from activation to NEBD of the second mitosis, and the time from NEBD to anaphase during the second mitosis of double-sized embryos shown in Fig. 3E and F. Statistical analysis was performed using the Mann-Whitney U test to compare the two groups.

**Figure S4.**
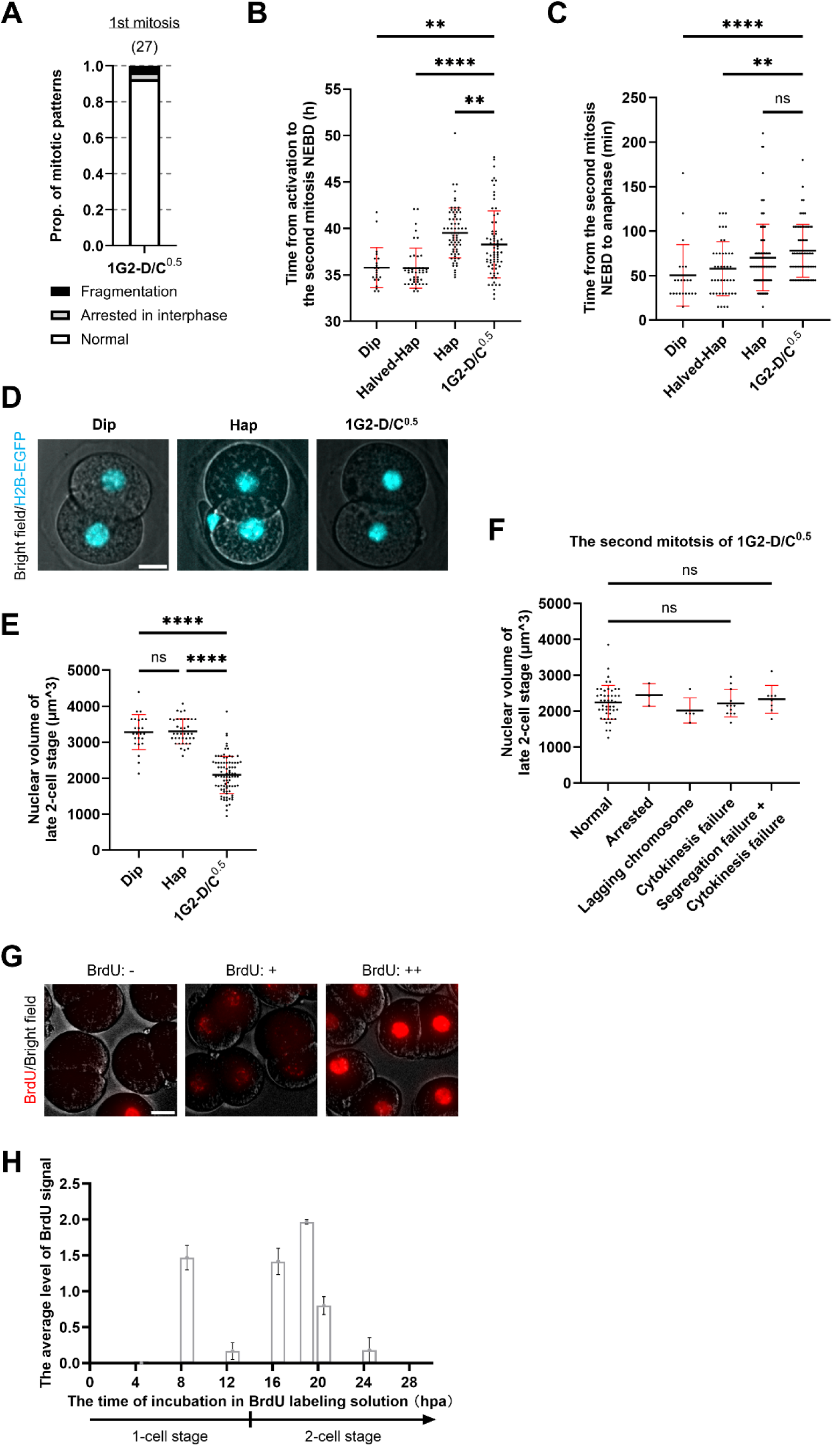
Embryos with a halved D/C ratio at the G2 phase of the 1-cell stage exhibit a prolonged 2- cell stage and reduced nuclear size compared to diploids. Relates to Figure 4. (A) Proportions of first mitosis conditions observed through time-lapse imaging (samples from three independent experiments). See related Video S4. (B and C) Scatterplots showing the time from activation to NEBD of the second mitosis, and the time from NEBD to anaphase during the second mitosis in 1G2-D/C^0.5^. The data used for comparison with the 1G2- D/C^0.5^ group is identical to Fig, S2A and B. (D) Representative still images from time-lapse imaging at the late G2 phase of the 2-cell stage in three types of embryo models. Embryos express histone-H2B-EGFP (cyan, z-projected). (E) Scatterplots showing the nuclear volume of three embryo models at the late G2 phase of the 2-cell stage. Nuclear volume was calculated by manually outlining nuclei that appeared approximately circular and assuming them to be spheres. Statistical analysis was performed using the One-way ANOVA to confirm overall significant differences among groups, followed by pairwise comparisons using Dunn’s multiple comparisons test to account for multiple comparisons. (F) Scatterplots showing the nuclear volume of 1G2-D/C^0.5^. The data are the same as those presented in E but classified based on the status of the second mitosis. Statistical analysis was performed using the 2-way ANOVA to confirm overall significant differences among groups, followed by pairwise comparisons using Dunn’s multiple comparisons test to account for multiple comparisons. (G) Representative immunofluorescence images showing z-projected BrdU signals in Dip samples. From left to right: no BrdU signal detected in the nucleus (BrdU: -), uneven BrdU signals present in the nucleus (BrdU: +), and BrdU signals observed throughout the entire nucleus (BrdU: ++). (H) A quantified graph based on BrdU signal levels, where BrdU: - = 0, BrdU: + = 1, and BrdU: ++ = 2, plotted according to the average BrdU signal at each time point. This was used as a reference for cell cycle assessment in enucleation experiments. Scale bars = 20 μm, sample sizes are indicated in parentheses above each bar. Note: The video files corresponding to the main figures are supplementary materials and are not included in this preprint. They are available from the authors upon reasonable request.

**Video S1. Haploid embryos exhibit mitotic abnormalities from the second mitosis onward, related to Figure 2**.

Representative embryos expressing histone-H2B-EGFP. Timestamps indicate hpa (hour:min) and scale bars are 20 μm.

**Video S2. Spindle structures within defective blastomeres were also aberrant during the second mitosis, related to Figure S2.**

Representative embryos injected with mRNA of mRFP1-tagged histone-H2B and EGFP-tagged α-tubulin.

Timestamps indicate the time from NEBD (in minutes) for the blastomeres indicated by the white arrow and scale bars are 20 μm.

**Video S3. Diploid embryos with doubled cytoplasmic volume exhibit mitotic abnormalities from the second mitosis onward, related to Figure 3**.

Representative double-sized embryos expressing histone-H2B-EGFP. Timestamps indicate hpa (hour:min) and scale bars are 20 μm.

**Video S4. The first mitosis of 1G2-D/C^0.5^ proceeds without abnormalities, related to Figure 4 and Figure S4.**

Representative 1G2-D/C^0.5^ expressing histone-H2B-EGFP. Timestamps indicate hpa (hour:min) and scale bars are 20 μm.

**Video S5. The second mitosis of 1G2-D/C^0.5^ exhibits mitotic abnormalities, related to Figure 4**.

Representative 1G2-D/C^0.5^ expressing histone-H2B-EGFP. Timestamps indicate hpa (hour:min) and scale bars are 20 μm.

**Video S6. The second mitosis of 2S-D/C^0.5^ exhibits mitotic abnormalities, related to Figure 5**.

Representative 2S-D/C^0.5^ expressing histone-H2B-EGFP. Timestamps indicate hpa (hour:min) and scale bars are 20 μm.

## Notes

### Competing Interest Statement

The authors have declared no competing interest.

